# A deep learning approach for improved detection of homologous recombination deficiency from shallow genomic profiles

**DOI:** 10.1101/2022.07.06.498851

**Authors:** Gregoire Andre, Tommaso Coletta, Christian Pozzorini, Ana C. Marques, Jonathan Bieler, Rieke Kempfer, Chloe Chong, Alexandra Saitta, Ewan Smith, Morgane Macheret, Adrian Janiszewski, Ximena Bonilla, Jaume Bonet, Hugo Santos-Silva, Magdalena Postl, Lisa Wozelka-Oltjan, Nils Arrigo, Adrian Willig, Christoph Grimm, Leonhard Müllauer, Zhenyu Xu

**Affiliations:** SOPHiA GENETICS SA, Rue du Centre 172, CH-1025 Saint Sulpice, Switzerland; Division of General Gynecology and Gynecologic Oncology, Gynecologic Cancer Unit, Comprehensive Cancer Center, Medical University of Vienna, Austria; Department of Pathology, Medical University of Vienna, Währinger Gürtel 18-20, A-1090 Vienna, Austria

## Abstract

Homologous Recombination Deficiency (HRD) is a predictive biomarker of poly-ADP ribose polymerase 1 inhibitors (PARPi) response. Most HRD detection methods are based on genome wide enumeration of scarring events and require deep genome sequence profiles (> 30x). The cost and workflow-specific biases introduced by these genome profiling methods currently limits clinical adoption of HRD testing.

We introduce the Genomic Integrity Index (GII), a Convolutional Neuronal Network, that leverages features from low pass (1x) Whole Genome Sequencing data to distinguish HRD positive and negative samples. In a cohort of 230 ovarian and breast cancer, we found GII supports accurate stratification of samples yielding results that are highly concordant with state-of-the-art HRD detection methods (0.865<AUC<0.996) which require 50x deeper coverage.

We conclude that the deep learning framework supporting GII allows accurate detection of HRD from shallow genome profiles, reducing biases and data generation costs making it uniquely suited for clinical applications.

## Introduction

Dysregulation of the complex pathway that safeguards genome integrity and prevents DNA damage is a common hallmark of cancer ^1^. This tumorigenesis-predisposing factor can also be exploited to induce cancer cell death ^2^. This is illustrated well by the response of Homologous Recombination Deficient (HRD) tumors to Poly-ADP ribose polymerase 1 inhibitors (PARPi) ^3^. PARPi induce the formation of DNA double-strand breaks (DSB) causing cell death in tumors with defects in the Homologous Recombination Repair (HRR) pathway that are unable to properly repair this type of DNA damage ^4^.

The diversity of somatic and germline loss-of-function events in HRR genes, including *BRCA1* and *2* ^5 6 7^, identified in PARPi responders ^8^, make patient stratification based on genotyping challenging. This is because mutations in several HRR-related genes have been shown to cause HRD but the impact on these genetic lesions in genome integrity and PARPi response is hard to predict ^3^ and can be tissue specific ^9^. Furthermore, epigenetic inactivation of HRR genes, which is not captured in genomic DNA sequencing, can also cause HRD, further limiting the sensitivity of methods that rely on detection of deleterious variants in HRR genes ^3^.

Defects in DNA repair pathways yield specific mutational signatures or genomic scars ^10^ which can be used as biomarkers. Specifically, the genome of HRD cells is characterized by the presence of small (<50 nucleotide) deletions flanked by regions of microhomology^11–13^. This pattern is similar to mutational signatures 3 and 8, which are enriched in the genome of HRD tumors ^14^. In addition, loss of HRR results in increased frequency of large-scale genomic rearrangements, including loss of heterozygosity (LOH) ^15^, Telomeric-Allelic Imbalance (TAI)^16^ and Large-Scale State Transitions (LST)^17^. Several methods use these features alone, or in combination, to identify HRD tumors ^9^. Methods that detect the presence of mutational signature alone^14 18^, or in combination with the presence of large-scale rearrangements, such as HRDetect ^19^, are amongst the most accurate in identifying HRD positive tumors. However, identification of cancer-related mutational signature with high sensitivity requires high coverage (>30x) whole-genome sequencing (WGS) data from tumor-normal pairs which is costly and hard to implement, and therefore not commonly used for cancer patient management.

The combined number of LOH, LST and TAI events detected in the tumor genome, Genomic Instability Score (GIS) ^20^ can be used to identify PARPi-responders with high confidence from tumor only data ^21^. However, detection of LOH and TAI requires Variant Allele Frequency (VAF), only accessible from deep genomic profiling data, which is uncommon in the clinical practice. Alternative methods which rely on detection of copy number changes, including LST events, from WGS at low sequencing depth (~1x) can also be used to predict tumor HRD status ^22, 23^ and provide cost-effective and easy to implement alternative for HRD detection. However, the power of methods that solely rely on this genomic scar to identify HRD samples is limited ^23^, and the full potential of low pass WGS (lpWGS) in HRD classification is unlikely to be unlocked using approaches that rely on establishing the frequency of biomarker events.

The genomic scars associated with HRD are expected to impact sequencing coverage genome wide. In addition, certain regions of the genome are more frequently impacted by DSB ^24^, raising the possibility that HRD-related scars are not homogenously distributed across the genome. Deep learning methods can be used to leverage the impact on the coverage profiles from lpWGS data of well-established or yet unexplored genomic scars induced by HRR loss-of-function to classify patients based on their HRD status.

Indeed, the ability of this class of algorithms in general ^25^, and Convolutional Neuronal Networks (CNN) in particular, to learn data-encoded patterns has supported unprecedented advances in many fields, including image processing ^26, 27^. Inspired by the visual cortex of the brain, CNN take an input image through a series of connected layers of convolution, data reduction filters and typically output a class label ^27^. The convolution filters are learnt from sample data during the CNN model training, making the performance of this class of methods extremely reliant on large volume data ^27^. This reliance on large datasets, which are often not available for the development of genomic analysis tools, has limited the use of CNN for this type of applications. Similar challenges in the development of clinical image analysis motivated the development of approaches, such as data augmentation, to *in-silico* increase in size and diversity of training datasets. Successful data augmentation strategies preserve the key properties of the augmented dataset and result in artificial data which cannot be distinguished from the original dataset ^28^. Data augmentation has been used to increase the quality and quantity of the training data for a variety of image classification tasks in molecular and cell biology ^29 30^ and biomedicine ^31^, but its use in genomics has so far been limited.

Here we describe the development of a CNN-based method that leverages the differences between the coverage image plots obtained from lpWGS data of HRD positive and negative samples to compute a Genomic Integrity Index (GII). We developed a novel data augmentation strategy to train the GII algorithm to overcome the lack of suitably large datasets. We validated the GII using lpWGS data of breast and ovarian cancer samples and showed it supports HRD detection, with accuracies similar to those obtained using methods that rely on data types less frequent in the clinical setting due to cost or implementation, making the GII a promising alternative for HRD detection in the clinics.

## Results

### The distinct lpWGS genome coverage profiles of HRD positive and negative samples

To simulate lpWGS (~1x) data, we down-sampled high coverage (~30-40x) WGS data from 274 FF breast cancer samples^19^ to a uniform depth of 10 million mapped paired-end reads (fragments). We computed the CG-normalized smoothed read coverage profiles from the simulated lpWGS libraries for these samples. According to HRDetect ^19^ that was considered, unless otherwise stated as ground-truth, 100 out of the 274 samples in the dataset were HRD positive. The remaining samples were classified as HRD negative.

We observed that the normalized coverage profiles of samples labelled as HRD positive, including those with somatic or germline loss-of-function mutation in *BRCA1* or *BRCA2*, were distinct from those of the remaining samples in the same cohort (Figure 1). Specifically, the genome instability resulting from HRD leads to visible differences in sequencing coverage plots, including distinctive and relatively frequent sharp differences in the depth of coverage of contiguous regions across the genome of HRD positive samples. In addition, whereas differences in coverage depth are apparent throughout HRD positive genomes, they are relatively more frequent in certain regions, consistent with the consequences of the genome instability caused by HRD being heterogeneous across the genome (Figure 1).

**Figure 1.**
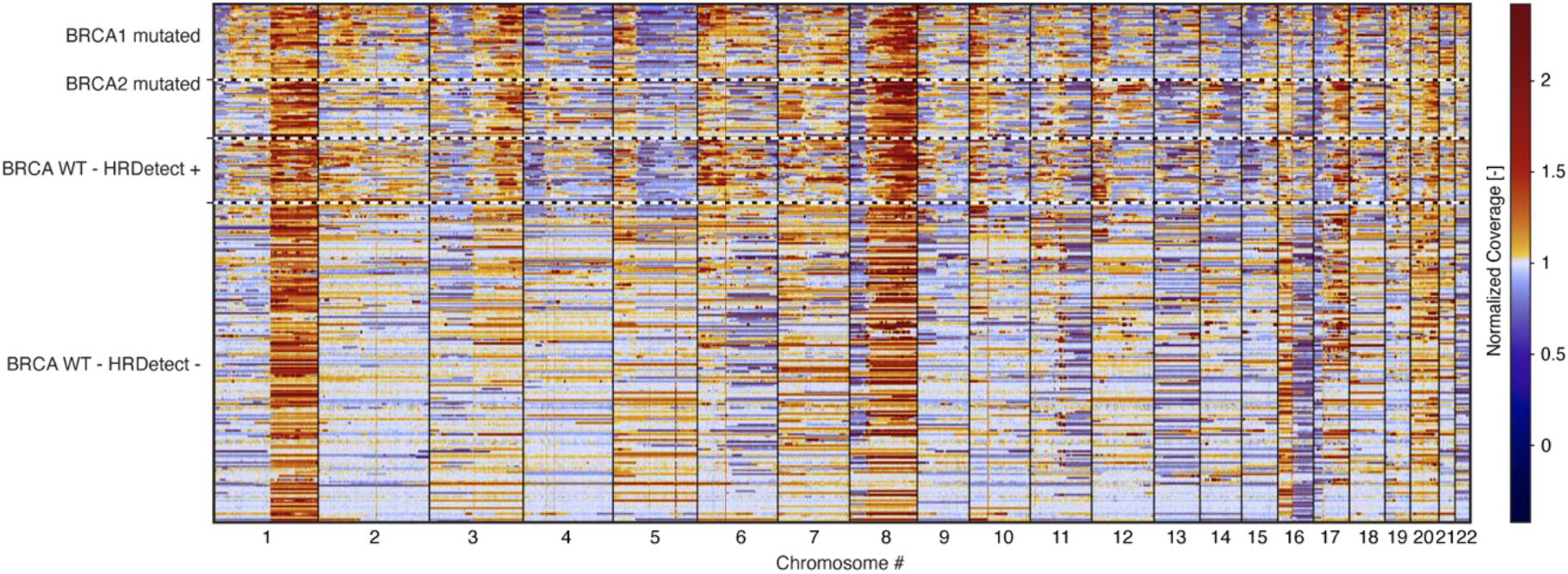
The normalized coverage plots of HRD positive and negative samples are distinct. Heatmap representation of the normalized coverage represented in bins of 100 kb lpWGS data for 274 FF breast cancer samples (x-axis). Sex chromosomes are excluded from this analysis. Samples are ordered from top to bottom, as following: *BRCA1* mutated; *BRCA2* mutated; *BRCA* wild-type reported as HRD positive by HRDetect ^19^ (sorted by decreasing HRDetect Score, HRDetect +); and *BRCA* wild-type (WT) reported as HRD negative by HRDetect ^19^ (sorted by decreasing HRDetect Score, HRDetect −). Bins are colored based on their normalized coverage relative to the mean coverage of the sample (set to 1, white). Color scale is depicted on the right.

### Development of a Convolutional Neuronal Network model for HRD detection

We reasoned that a machine learning algorithm could leverage the differences caused by genome instability on coverage plots from lpWGS data to classify HRD positive and negative samples.

The development of this type of algorithm for genomic analysis of clinical biomarkers is often limited by the relatively small number of samples available for training. For this reason, we turned our attention to image classification algorithms, in particular CNN, that require fewer parameters than other machine learning methods, thus limiting the risk of overfitting ^32^.

In addition, and to overcome the challenge posed by the limited number of samples available for training, we implemented a data augmentation strategy to increase the number and diversity of samples in our training dataset. Augmented training data was generated by randomly sampling chromosomes from a subset of the original data ^19^ (173 breast cancer samples; 61 HRD positive of which 47 BRCA mutated, 35% and 27% respectively) (Figure 2A). To generate one data augmented sample, we combined chromosomes from a set of randomly selected samples with the same HRD status (Figure 2A). We accounted for the impact of sample specific differences in purity and ploidy by normalizing the purity and ploidy ratio (purity and ploidy ratio=purity_sample_/ploidy_sample_) of all samples used to generate one data augmented sample (detailed in the methods section, Figure 2B & 2C). Data augmented samples (Figure 2D) were obtained by combining chromosomes from samples with normalized purity/ploidy, ensuring the amplitude of coverage differences observed for a given ploidy was constant across chromosomes. This approach introduced biases in the purity/ploidy ratios of data augmented samples which tend to be lower than what was observed in the original samples (Figure 2E). To account for this bias, we applied a Metropolis-Hastings and Gibbs sampling method to ensure that the purity/ploidy distribution of the 3760 retained data augmented training samples matched that of the original samples (detailed in the methods section, Figure 2F), ensuring that after data augmentation the properties of the training data reflect those of real data.

**Figure 2.**
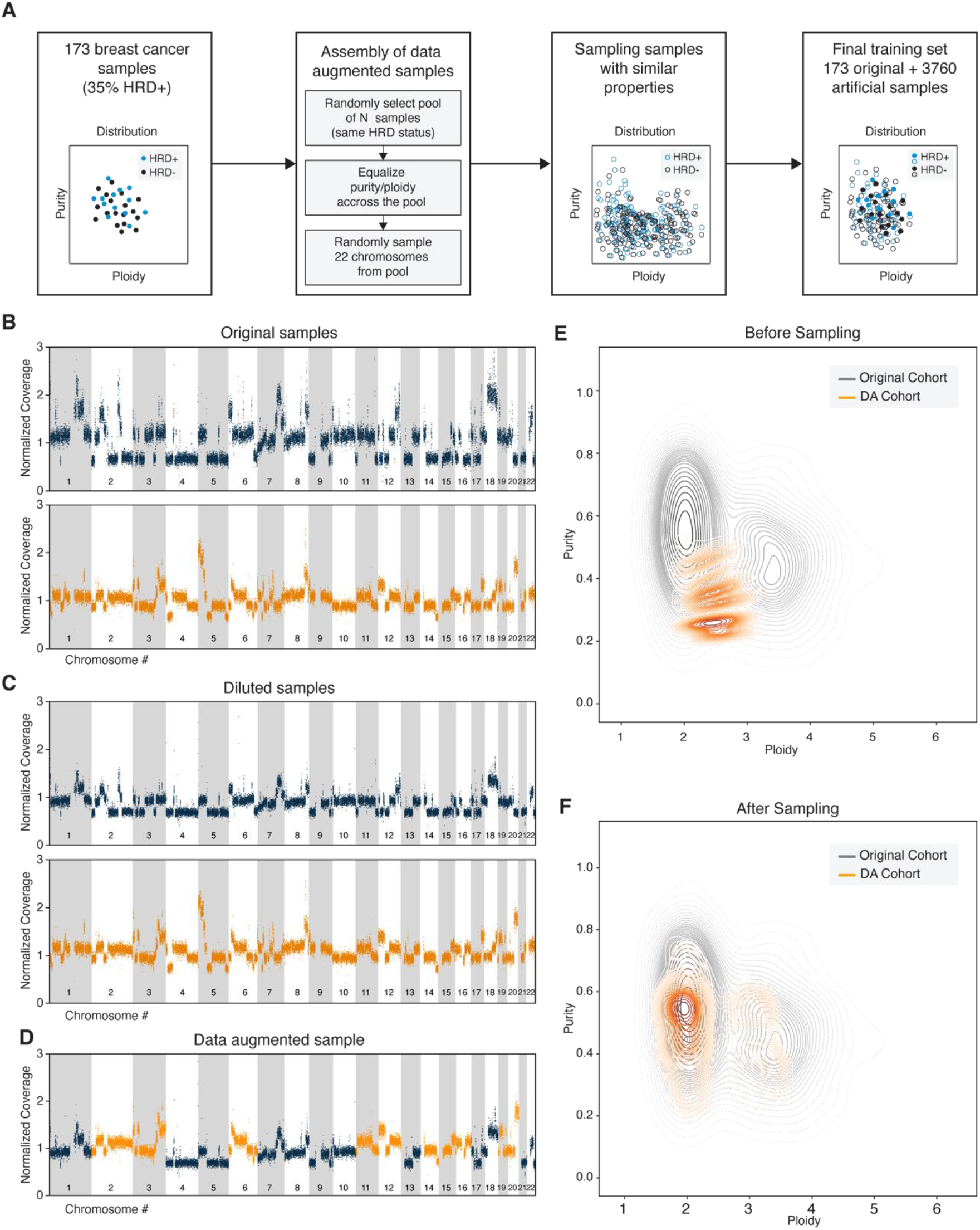
Using data augmentation to increase the size and diversity of the training set. (A) Schematic representation of data augmentation strategy. Each dot represents the purity (y-axis) and ploidy (x-axis) of one HRD positive (+, turquoise) and negative (-, navy) samples in the original (closed circles) or data augmented (open circles) dataset. Normalized coverage profile for 2 exemplary samples (B) before and (C) after *in-silico* decreasing of sample(s) purity. Decreasing purity ensures the purity and ploidy ratio matches that of the sample(s) with the smallest ratio observed in the set of samples that make up the exemplary (D) data augmented sample. Normalized coverage (y-axis, ranging from 0-3) is depicted for each genomic bin (x-axis, autosomes are ordered left to right from 1-22). Each sample is represented by a different color. Data originating from sample A and B are colored navy and orange throughout. Sample A has purity of 0.80 and ploidy of 1.84. Sample B has purity of 0.49 and ploidy of 2.29. Illustration using HRD positive samples of the difference between the distribution of sample purity (y-axis) and ploidy (x-axis) in the original (grey) and data augmented (orange) data set before (E) and after (F) Metropolis-Hastings and Gibbs sampling.

We note that the normalized coverage plots from data augmented samples (Supplementary Figure 1) are similar to those obtained for the original samples (Figure 1). We considered 3933 samples (173 patient-derived 3760 data augmented samples) to train, using a supervised learning framework, a CNN model that uses lpWGS sequencing information to classify samples according to the HRD status reported by HRDetect ^19^. It was previously noted that genome instability caused by HRD observed along the genome is not homogenous. For example, large scale transitions near chromosome centromeres correlate poorly with genome instability caused by HRD ^17^. We hypothesized that spatially arranging data from autosomes would facilitate identification by the CNN model of this and other regional differences in genome instability caused by HRD. We created 2 dimensional heatmaps where each row corresponds to one autosome, ordered from 1-22, and each column represents a genome bin of 3Mbp aligned with respect to their respective centromere (Figure 3A). Throughout the manuscript we refer to these heatmaps as smoothed coverage plots.

**Figure 3.**
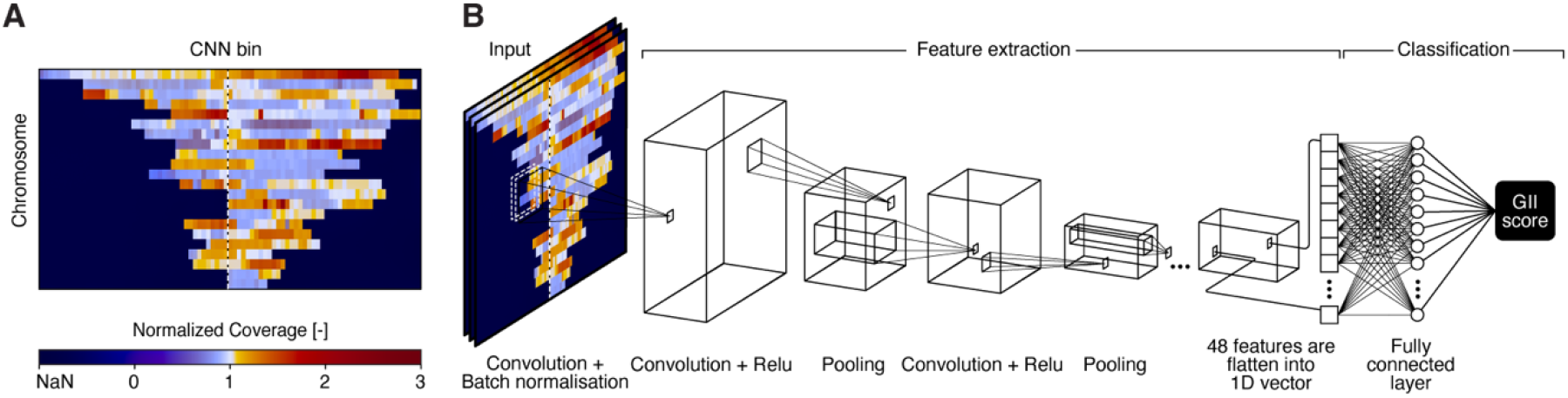
GII was developed to predict HRD status using spatially organized coverage profiles from lpWGS. (A) Example of GII input image. Smoothed normalized coverage across ~3Mbp bins (columns) for all autosomes (rows) aligned with respect to their centromere (vertical dashed line). Bins are colored (blue-white-red scale) based on their normalized coverage relative to the mean coverage of the sample (set to 1, white). Color scale is depicted on the bottom. (B) Schematic of GII architecture. Features in input image are extracted through series of convolution and pooling operations. The vector containing the 48 extracted features is provided to a set of fully connected layers trained to predict the samples GII status.

The CNN model that takes smoothed coverage plots was trained following a 5-fold cross validation procedure. The CNN contains 3 convolution blocks that output a scalar, maximizing the differences between HRD positive and negative samples in the training set (Figure 3B, Materials and Methods).

### Validation of GII in Breast Cancer

To evaluate the analytical performance of GII, we considered WGS data for 101 FF breast cancer samples ^19^ not used to train the algorithm. We randomly down-sampled libraries to the equivalent of ~1x coverage and used GII to predict the HRD status of these samples based on the corresponding lpWGS smoothed coverage plots. We first compared the distribution of GII between samples from patients with and without mutations in *BRCA1/2*. Supporting GII’s ability to classify HRD positive and negative samples, the score distribution between BRCA mutated and wild-type samples was different (Figure 4A), with BRCA mutated samples displaying a higher score. To evaluate the performance of GII in distinguishing the two sample types and to compare it with other HRD detection methods, we determined the Area under the Curve (AUC) of the Receiver Operating Characteristics Curve (ROC) obtained when BRCA mutational status (Figure 4B) was considered as ground-truth (Materials and Methods, Figure 4B). All tested methods displayed relatively high concordance with BRCA mutational status (AUC>0.763, Figure 4B), with GII yielding the highest AUC (AUC=0.858) after HRDdetect, the best performing method (AUC=0.862).

**Figure 4.**
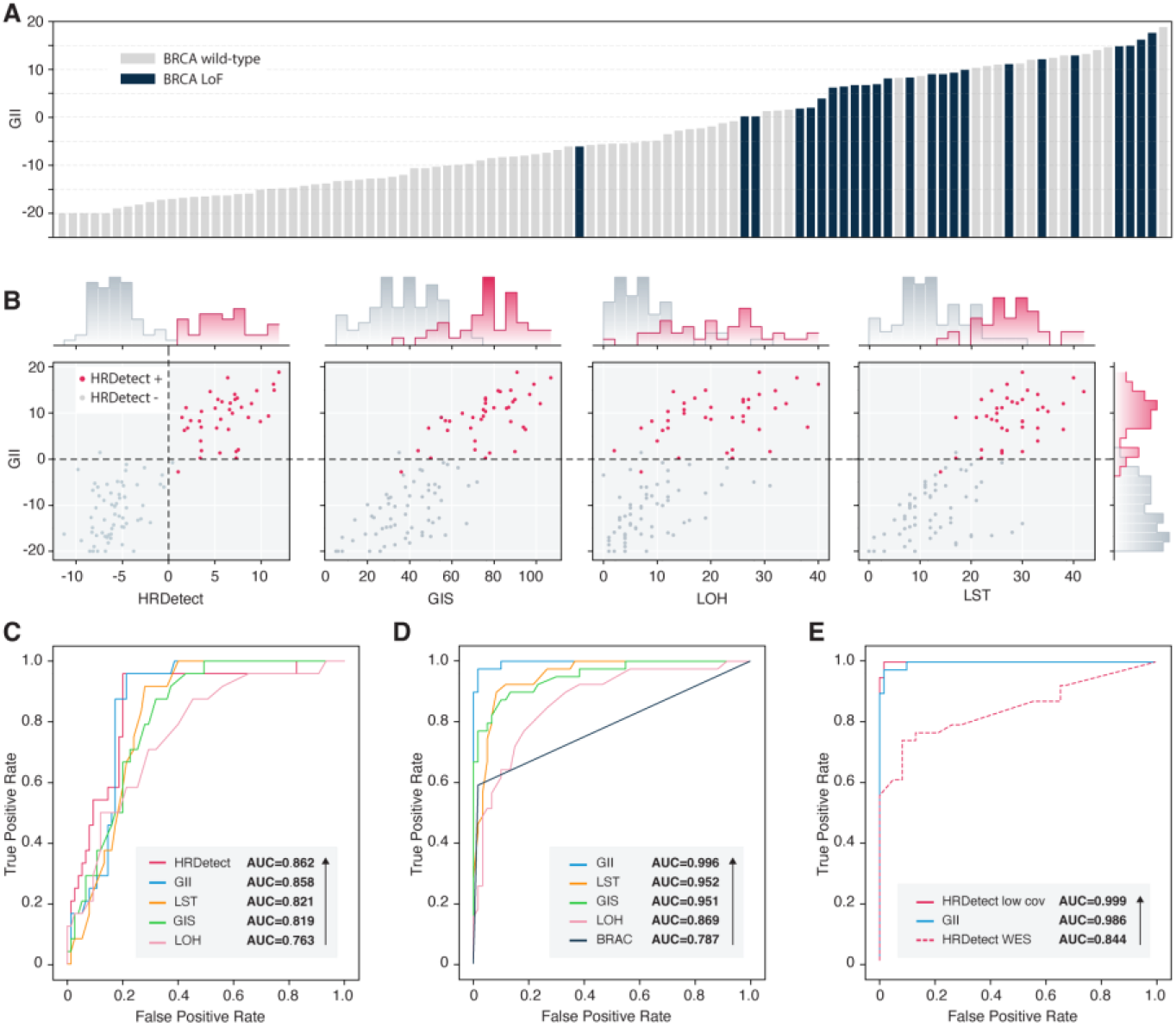
GII yields comparable results to tools that rely on high coverage datasets. (A) GII score (y-axis) obtained for different breast cancer samples (x-axis), ordered by GII score and colored according to their BRCA mutational status as BRCA wild-type (grey) and BRCA loss of function (LoF, navy). (B) GII score (y-axis) relative to the corresponding HRDetect, GIS, LOH and LST score (x-axis). Each point corresponds to the pre-sigmoid GII score for a breast cancer patient sample. HRD positive and negative samples according to HRDetect are colored in red and grey respectively. Histograms representing the distribution of the respective score were added to axes. (C) ROC curves for HRD classification obtained using HRDetect (red), GII (turquoise), LST (orange), GIS (green), LOH score (pink) using BRCA mutational status as ground-truth. AUC values are given for each method (inset). (D) ROC curves for HRD classification obtained using GII (turquoise), LST (orange), GIS (green), LOH score (pink) using HRD status reported by HRDetect as ground-truth. AUC values are given for each method (inset). (E) ROC curves for HRD classification obtained using HRDetect low coverage (red), GII (turquoise) and HRDetect WES (dashed red line), using HRD status reported by HRDetect as ground-truth. AUC values are given for each method (inset).

HRD can by caused by mutations or epimutations in other HRR genes and detection of BRCA loss of function mutations is less accurate in predicting HRD than methods that leverage on the genome instability resulting from HRD ^19^. We compared the distribution of GII score for samples with different HRD status according to HRDetect; the number of HRD biomarker events (LOH and LST score alone or combined with TAI GIS ^20^, Figure 4C). We found that GII obtained for samples classified as HRD+ or HRD-by these different methods was distinct, supporting the relevance of HRD classifications based on GII score (Figure 4C).

As expected, given HRDetect labels were used to train GII, the classification by GII within this cohort was most similar to this tool (AUC=0.996), followed by LST and GIS (AUC=0.951 and AUC=0.952, respectively) (Figure 4D). The concordance with other methods which explore individual features previously associated with HRD, including the number of LST events computed using lpWGS data^23^, was relatively low. This finding supports that GII leverages other HRD relevant features from lpWGS.

It was previously shown that the analytical performance of HRDetect correlates with sequencing depth, detecting higher accuracy when sequencing coverage is relatively high (~30x) ^19^. Since GII takes advantage of lpWGS data (~1x), we wanted to compare its performance to HRDetect results obtained using data at lower coverage (~10x) or exon sequencing data, available from the original study ^19^. Despite relying on lpWGS data, the results of GII are most concordant with HRDetect classification rather than HRDetect results based on exon sequencing (AUC=0.844, Figure 4E) data. Despite relying on 10-times fewer coverage with GII, the concordance classifications yielded by analysis of HRDetect using coverage of 10x (AUC=0.999) was similar to what was obtained using GII (AUC=0.996). This analysis supports that the information leveraged by GII from lpWGS data allows to distinguish HRD positive and negative tumors in breast cancer with accuracies matching those of well-established methods.

### Validation of GII on FF ovarian cancer samples

Loss of function of the HRR pathway has prognostic value in other cancer types ^33, 34^. To gain initial insights into the relevance of GII in other tumor types, we evaluated WGS data ^19^ for 63 FF ovarian cancer samples in which *BRCA1/2* loss-of-function and HRD are also very common ^33, 34^. Similar to the analysis in breast cancer samples, GII distribution was different between BRCA mutated and BRCA wild-type ovarian cancer samples (Figure 5A). The value of GII in classification of these samples was also apparent by the high AUC associated with using GII to predict samples’ *BRCA1/2* status (AUC=0.935, Figure 5B). GII concordance was lower than for HRDetect (AUC=0.982, Figure 5B) but higher than what can be obtained using LST and LOH, also able to be calculated from lpWGS (AUC>0.905, Figure 5B). The confidence on differences in AUC between methods was limited by the relatively small cohort size available for testing.

**Figure 5.**
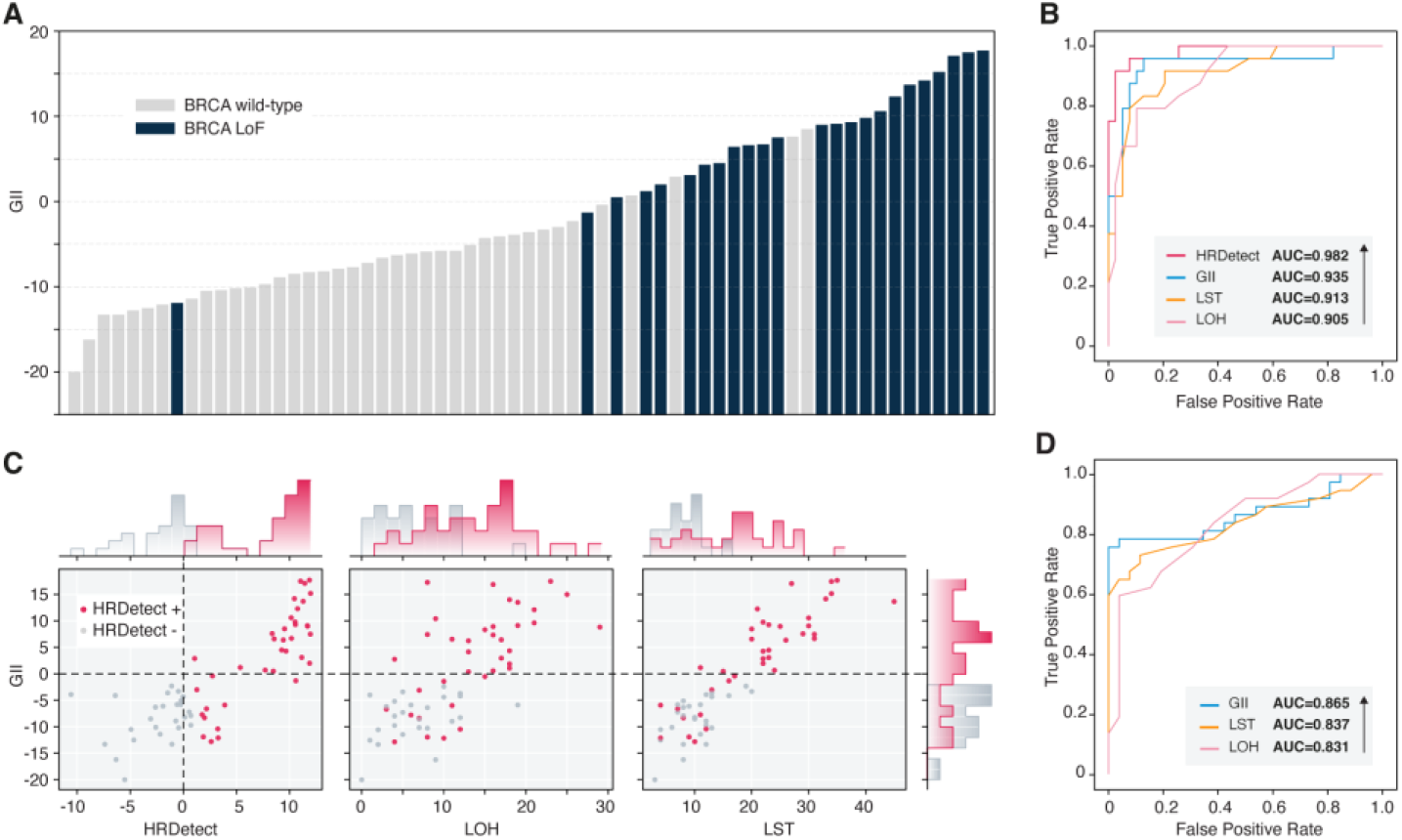
The GII accurately predicts *BRCA1/2* and HRD status in ovarian cancer. (A) GII score (y-axis) obtained for different ovarian cancer samples (x-axis), ordered by GII score and colored according to their BRCA mutational status as BRCA wild-type (grey) and BRCA loss of function (LoF, navy). (B) ROC curves for HRD classification obtained using HRDetect (red), GII (turquoise), LST (orange) and LOH score (pink) using BRCA mutational status as ground-truth. AUC values are given for each method (inset). (C) GII score (y-axis) relative to the corresponding HRDetect, LOH score and LST (x-axis). Each point corresponds to the pre-sigmoid GII score for an ovarian cancer patient sample. HRD positive and negative samples according to HRDetect are colored in red and grey respectively. Histograms for the scores were added to axes. (D) ROC curves for HRD classification obtained using GII (turquoise), LST (orange) and LOH score (pink) using HRD status reported by HRDetect as ground-truth. AUC values are given for each method (inset).

Next, we compared the results obtained by GII to the alternative classification results available for these samples. The GII score distribution for ovarian cancer samples, classified by HRDetect, LOH and LST score as HRD positive or negative was different (Figure 5C), supporting the relevance of this metric for sample stratification. Whereas still relatively high (AUC=0.865, Figure 5D), the overall concordance between HRDetect and GII in this cohort was lower than what was seen in breast cancer cohort (Figure 4D) and what was obtained when BRCA status in ovarian cancer samples was used as ground-truth (Figure 5B). This is consistent with the presence of tissue specific differences in the features used by machine learning algorithms to stratify samples and are likely to impact HRD calls. For example, the analytical performance for HRDetect in non-breast cancers samples was reportedly lower, likely a consequence of the use of mutational signatures which are impacted by the mutational patterns in the tissue of origin ^19^. Of note, in the absence of information on PARPi response for ovarian cancer patients in the cohort, the discrepancies between methods cannot be unequivocally interpreted.

Nonetheless, GII results are more highly concordant with HRD calls than are the results available for these samples obtained using the alternative state of the art method, i.e. LOH (AUC-0.831, Figure 5D) and LST, which can be estimated from lpWGS (x1) (AUC 0.837 Figure 5D).

Based on the data, we conclude that GII predicts HRD status in non-breast cancer samples with high accuracy.

### Validation of GII in Clinical samples

Clinical samples are often preserved as formalin-fixed, paraffin embedded (FFPE) blocks to ensure cost-effective long-term storage. Relative to their FF counterparts, FFPE samples have lower DNA quality, which is known to make detection of a wide range of genetic signatures challenging ^35^. To gain initial insights into the performance of GII on FFPE samples, we generated lpWGS data (~1x) from DNA of 66 FFPE ovarian cancer biopsies collected between 2015-2019 (Supplementary Table S1). As expected, the fraction of high molecular weight DNA, measured by using the DNA Quality Number (DQN), tends to decrease with increasing sample age (Figure 6A). Samples with < 30% of DNA fragments above 300 bp (DQN<3, 1 sample) were excluded from the analysis. The tumor content of samples in the cohort, measured by estimating the percentage of tumor cells on hematoxylin and eosin stained slides, varied between 2% and 95% (Figure 6B). Since limited tumor DNA will dilute the signal from genome instability signatures, we restricted our analysis to samples with at least 30% tumor content which is in line with what is recommended for other methods that also leverage genome instability to detect HRD ^36^.

**Figure 6.**
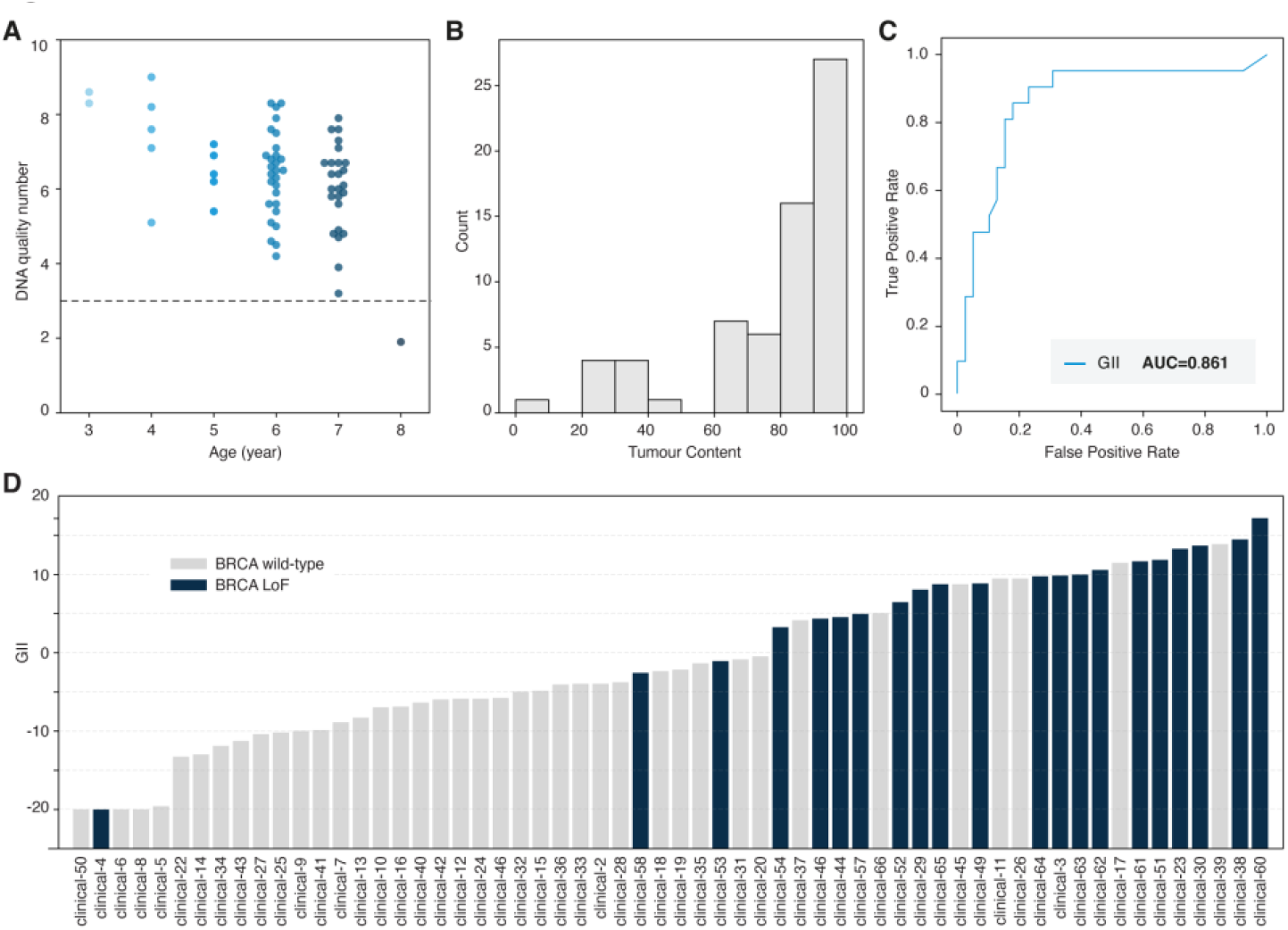
GII can be used to analyze FFPE samples. (A) DNA quality number (y-axis) for ovarian cancer samples as a function of the sample age (in years) at the time of DNA extraction (x-axis). Each point represents a sample. The horizontal line (DQN=3) indicates the minimal DQN of samples considered for GII analysis. (B) Histogram of clinical sample tumor content. (C) ROC curves for HRD classification obtained using GII (turquoise), using *BRCA1/2* status as ground-truth. AUC values are given for each method (inset). (D) GII score (y-axis) obtained for different ovarian cancer samples, ordered by GII score and colored according to their BRCA mutational status as BRCA wild-type (grey) and BRCA loss of function (LoF, navy).

We used targeted capture followed by sequencing to determine the mutational status of *BRCA1/2* that was used as ground-truth when assessing GII’s analytical performance in this cohort. We used BRCA exchange ^37^ to annotate variants in *BRCA1/2* as BRCA wild-type or loss of function (LoF, Materials and Methods). The results obtained based on lpWGS data for *BRCA1/2* wild-type and mutated samples supports that GII can distinguish HRD positive and negative samples (Figure 6C-D). Specifically, the GII score for samples with BRCA loss of function mutations was higher than what was obtained for BRCA wild-type samples (Figure 6D. In this cohort, the concordance between BRCA status and GII was high (AUC=0.861, Figure 6C. We observed 12 *BRCA1/2* wild type samples with a score > 95% quantile of BRCA loss of function samples (Figure 6C). This is compatible with HRD positive BRCA wild-type samples carrying similar genomic scars as their BRCA LoF counterparts ^10^. The GII score for one sample (Clinical-4) with a *BRCA1* missense variant annotated by BRCA exchange as pathogenic (NM_007294.3:c.181T>G) was low (GII = −20) and similar to the score obtained for most BRCA wild-type samples (Figure 6D). In contrast to what we found for the remaining *BRCA1/2* LoF samples (Supplementary Figure 2A), the normalized coverage profile of Clinical-4 sample did not display the abrupt changes in coverage characteristic from HRD scarring (Supplementary Figure 2B). In light of the high tumor content of this sample (~80%, Supplementary Table 1), the lack of abrupt changes in the normalized coverage profile was indicative of the lack of copy number changes in the sample. The variant fraction of the *BRCA1* pathogenic variant in Clinical-4 sample was 45%. Together, these observations suggest that the variant of interest only affects one of the 2 gene copies present in the tumor. In line with what has been observed by others ^19^, this result indicates that monoallelic inactivation of *BRCA1/2* may not result HRR deficiency and associated scarring.

Whereas analysis of a larger panel of clinical samples will be necessary to assess whether GII can be used for diagnostic purposes and to the determine GII threshold that supports accurate HRD classification, the results obtained on FFPE samples support the use of GII in clinical samples.

## Discussion

Cancer patients with HRD tumors respond favorably to PARPi treatment ^8^. Given the diversity of genetic and epigenetic events underlying HRD ^3^, methods which rely on detection of genetic lesions on a subset of well-defined genes can fail to identify patients that could benefit from PARPi treatment. Approaches that rely on detecting the consequences of loss of HRR function allow for patient stratification in a more precise manner. One consequence of cells’ inability to repair DSB, is the accumulation of small- and large-scale genomic lesions, a phenomenon known as genome instability ^3^. The increase in sensitivity of methods that explore biomarker events is a testimony of the added value of leveraging the consequences rather than the causes of HRD, bring to patient stratification^2^. However, detection of some of these scars, including LOH or TAI, require deep genomic profiling (>30x coverage) to support variant calling. The bias and the cost associated with methods that support the generation of deep genome profiles currently limit HRD testing in the clinical setting. Our aim was to develop a sample stratification method that explores the consequences of HRD on genome stability and can be deployed in clinical routine.

To ensure this method can be deployed in the clinical setting, we decided to rely on the use of lpWGS data, which is both cost-effective and easy to implement. The GII leverages the impact on genomic lesions based on lpWGS coverage plots to stratify patients based on their genome instability. The deep learning framework allows GII to go beyond prior knowledge and expert design, and to identify features that optimally allow for the distinction of HRD positive and negative samples from low-coverage data.

We used publicly available data to benchmark GII against other well-established methods of HRD detection. GII results are most similar to those of HRDetect, which relies on WGS (~10-50x coverage) to enumerate large scale events and genomic signatures characteristic of HRD. Concordance was also high with GIS, which integrates the number of HRD biomarker events ^20^. Both methods rely on the quantification of features which are only accessible from high coverage data. Similar classifications obtained from GII using data sets that have at least 10x higher sequencing depth indicate that this approach is an alternative and cost-effective approach for HRD detection. In addition, due to the fact it supports the use of WGS data at limited coverage, GII can, in principle, also be used for HRD detection from data generated from single cells, including circulating tumor cells.

Similar to the other machine learning method (i.e., HRDetect), the accuracy of GII is higher in breast cancer. This is likely the result of the model having been trained using sequencing data from this type of cancer patients, and it is indicative of tissue-specific differences in the strength of the features used by machine learning methods for sample classification. The availability of larger pan-cancer cohorts with HRD status will allow to account for this limitation, for example by using methods such as transfer learning, supporting the retraining of the final CNN layers and appraise tissue specific factors. Despite this limitation, the accuracy of the GII in non-breast cancer samples remains high (AUC=0.94) and comparable to that of methods which rely on 50x higher sequencing depth (AUC=0.98).

Furthermore, our analysis of a cohort of FFPE ovarian cancer samples supports that GII can distinguish BRCA mutated and wild type samples with high accuracy (AUC=0.86). Analysis of a larger clinical cohort is now needed to establish GII score threshold which allows optimal stratification of HRD samples and to determine what sample features should be considered to support decentralized use of GII in HRD testing.

The main limitation of GII and other methods, which rely on the detection of the genomic impact of HRD, is that genome instability scars accumulate over time, resulting in the possibility of false negatives due to insufficient time since the loss of HRR function. Similar to other methods, integration of GII with results of variant calling analysis in HRR genes should limit the impact of this constraint.

In summary, GII is a deep learning method which allows accurate stratification of HRD samples, including FFPE, based on lpWGS data. The accuracy of this method is comparable to that obtained using well-established methods, which typically require genomic data that is either more expensive or harder to obtain, making this a promising solution for HRD testing in the clinical setting.

## Methods

### DNA extraction

Tissue samples were analyzed from patients diagnosed with high-grade serous or endometroid grade 2 or 3 ovarian cancer between 2015 and 2019 in the Department of Pathology of the *General Hospital Vienna/Medical University Vienna*. The properties of the clinical samples are listed in Supplementary Table S1. The sample tumor content was obtained by estimating the percentage of tumor cells on hematoxylin and eosin-stained slides. DNA was extracted using the Promega Maxwell FFPE Plus DNA Kit. Briefly, depending on the tumor size, 1 to 5 FFPE sections were placed in a safe lock tube to which proteinase K and incubation buffer were added. Samples were incubated overnight on a thermal shaker set at 70°C and 1000 rounds per minute (RPM). Samples were then cooled down on ice. If a paraffin layer was visible, the supernatant was transferred into a fresh safe lock tube. This was followed by a lysis step and DNA extraction was performed automatically with the Maxwell® RSC Instrument (Promega) by running the program Maxwell® RSC FFPE Plus DNA Kit (AS1720). DNA was eluted in 60μl nuclease free water (NFW) and quantified using the Qubit® dsDNA HS Assay Kit (ref Q32851).

DNA quality was assessed by analyzing the fragment size distribution of the sample using the Agilent Fragment Analyzer. The DNA quality number (DQN) for each sample was determined as the fraction of DNA fragments larger than 300 bp using the DQN function of the Fragment Analyzer software, with the DQN threshold set to 300 bp.

### Library preparation and sequencing

Whole-genome and targeted libraries were prepared using SOPHiA GENETICS library preparation kit. First, 50 or 100 ng of artificial FFPE DNA input was end-repaired and A-tailed, followed by ligation to Illumina compatible adapters. Ligation products were purified using AMPure beads (Beckman Coulter) and further amplified by PCR for 8 cycles. Amplified libraries were cleaned up using AMPure beads (Beckman Coulter). The obtained libraries were used either at this step as WGS libraries or further enriched for targeted sequencing as following. Libraries were pooled and mixed with human Cot-1 DNA (Life Technologies) and xGen Universal Blockers-TS Mix oligos (Integrated DNA Technologies) and lyophilized. Pellets were resuspended in a hybridization mixture, denatured for 10 min at 95°C and incubated for 16 h at 65°C in the presence of biotinylated probes (xGEN Lockdown IDT®).

The probe panel spanned 157 Kb and covers a set of clinically relevant genes implicated in homologous recombination related (HRR) genes. Probe-hybridized library fragments were captured with Dynabeads M270 Streptavidin (Invitrogen) and then washed. The captured libraries were amplified by PCR for 15 cycles and cleaned up using AMPure beads (Beckman Coulter).

For a given sample, whole-genome and targeted libraries were added to flow cell in a 67%-33% ratio and sequenced to approximately 16M fragments per sample corresponding to ~1-2x whole genome coverage using NovaSeq S4 sequencing (Illumina) with 150 bp paired-end reads.

### Clinical sample sequencing data processing, variant calling and interpretation

Read alignment to the reference genome human genome (hg19), read filtering and variant calling were performed using the SOPHiA GENETICS proprietary analysis workflow. Briefly, after read mapping adaptors were trimmed, mispriming events were removed and read softclipped regions were realigned. Read fragments shorter than 21 bp were excluded, and the resulting alignment was used for variant calling using SOPHiA GENETICS pipeline and to prepare coverage plots (refer to preparation of coverage plot session).

To determine the BRCA status we annotated all *BRCA1* and *BRCA2* variants with BRCA Exchange (release available on the 13/12/2021). Variants reported in BRCA Exchange as either: ‘pathogenic’, ‘likely pathogenic’, ‘risk factor’, ‘probable pathogenic’, based on all, non-conflicting clinical significance reports, were annotated as ‘pathogenic’. Samples with at least one pathogenic *BRCA1* or *BRCA2* variant were classified as BRCA loss-of-function. The remaining samples were classified as BRCA wildtype.

### Processing of publicly available whole genome sequencing data

BAM files for WGS data for FF breast and ovarian cancer with BRCA-deficiency status were downloaded from Wellcome Trust Sanger Institute and the International Cancer Genome Consortium ICGC available at the European Genome-phenome Archive EGA (https://www.ebi.ac.uk/ega/studies/EGAS00001001178). This cohort was divided into training (173 breast cancer samples) and test set (101 and 63 breast cancer and ovarian cancer samples). HRD scores for these samples were obtained from ^19^.

### Preparation of Coverage plots

The human reference genome (hg19) was divided into contiguous non-overlapping intervals of 100kb (hereafter to as “bins”). Bins containing blacklisted regions, including regions known to introduce bias in coverage were excluded. These regions included: i) centromere and telomere positions as obtained from UCSC for hg19, ii) genome assembly gaps in the reference genome (N’s), iii) Regions with extreme GC content and high mappability, and iv) sex chromosomes. In the case of NGS data generated in a workflow combining both lpWGS and targeted sequencing, bins overlapping with enriched regions or regions highly homologous to enriched regions, were also discarded. The raw coverage count for each bin was obtained by counting the number of mapped pair-end reads (fragments) to the respective bin. Fragments mapping across two bins were assigned to the bin with the largest overlap. Only unambiguously mapped fragments were considered in the coverage calculation.

For TCGA data sets and to mimic lpWGS, data sets were down-sampled to 10 million pair-end reads (~1x coverage). The probability of observing *k* mapped fragments within a given genomic bin according to the multinomial distribution was considered : P(K=k) = n!/(k!(n-k)!) *p^k*(1-p)^(N-k), where *n* denotes the number of fragments after down-sampling and *p* is the prior probability that a fragment drawn will fall within the considered bin of the coverage profile defined as *p*=*C*_*bin*_/C_*to*t_, where *C*_*bin*_ and *C*_*tot*_ is the observed coverage for the bin considered and the sum of the raw coverage across all bins, respectively.

The raw coverage count of bins was normalized by dividing it by the sample’s mean coverage. Next, a lowess regression was fit to the raw coverage versus GC content of the bins. Corrected normalized coverage was obtained by dividing the normalized coverage by the result of the lowess fit. The normalized coverage data was further collapsed into an average 3Mbp target bin (size in the 2.5Mpb-3.5Mbp range) using the adaptive binning strategy and converted into a heatmap. The normalized coverage data for each sample was spatially arranged to form a 2D array of 87 bins * 22 autosomes. The coverage bins for 22 chromosomes were plotted as 22 rows from chr1 to chr 22, aligned with respect to their centromeric bins. For shorter chromosome arms, empty bins were filled by copying the value of nearest telomeric bin present in the same row.

### Data augmentation

173 breast cancer samples were considered in the training set (35% HRD positive, 27% *BRCA1/2* mutated) to generate data augmented (DA) dataset. To assemble one DA sample, 22 autosomes were randomly sampled from *N* original tumor samples with the same HRD status, where *N* is a number between 2 and 22 drawn from an exponential distribution with associated probability density function:

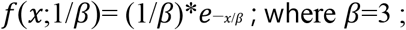

To account for biases in coverage plots introduced by differences in purity/ploidy ratio between samples, sample with the minimal purity/ploidy ratio (*m*) observed amongst the *N* samples used to assemble a given DA sample was determined. The other samples were *in-silico* diluted (decreased the sample purity) until their purity/ploidy ratio was equal *m*. This was achieved by mixing the normalized coverage profiles of the sample with the profile of one randomly selected normal sample. Next, DA augmented sample genomes were assembled by randomly selecting autosomes from the *N* samples after the purity/ploidy normalization step. Purity and ploidy information for each sample was provided by HRDetect authors for each sample ^19^. To ensure the distribution of the final augmented dataset in (purity, sample ploidy) space was representative of the distribution of the original breast cancer cohort, Metropolis-Hastings and Gibbs sampling was applied. The rejection sampling algorithm stopped when no new samples can be drawn after 100 000 iterations.

Spatially arranged normalized coverage plots for data augmented samples were obtained as described for the original tumor samples.

### Convolutional Neuronal Network

173 patient-derived and 3760 data augmented samples to train a CNN model (GII) were used to predict sample’s HRD status. The input to GII were smoothed coverage plots. 2-dimentional heatmap of the smoothed normalized coverage depth was obtained for the sample. Each row in the heatmap corresponds to one autosome, ordered from 1-22, and each column is a genome bin of 3Mbp (ordered from 5’ to 3’). Chromosomes were aligned with respect to their respective centromere.

The GII architecture included 3 convolution blocks (convolution, batch normalization and max-pooling) whose output was aggregated by an average pooling layer. The 48 features extracted by these convolution steps was flattened into 1D feature vector and passed to a fully connected layer with a single output node (whose output is a scalar) that was passed through a corresponding sigmoid activation function. Five CNN models with the described architecture and initiated using a different random seed were trained following a 5-fold cross-validation procedure. During the training step, Adam optimizer function^38^ was used to minimize the result of the binary-cross entropy loss function. To minimize the impact of model initialization, the final GII score was computed as the average of the GII scores obtained from the 5 CNN models.

### Performance evaluation

Algorithm performance was assessed using the Area Under the Curve (AUC) in the Receiving Operator Curves (ROC) using either BRCA mutational status or HRD classification according to HRDetect was considered as ground-truth using scikit-learn python library (v.0.24.0) ROC and AUC functions (default parameters). Except for GII and LST in FF ovarian cancer samples, all scores were obtained from the HRDetect publication ^19^. LST was obtained using scarHRD, using as input allele specific copy number segmentation profiles from FF ovarian cancer samples. To obtain these allele specific segmentations, Hidden Markov Model (HMM) was used which takes as input normalised WGS coverage profiles (100 kb) and variant fraction of likely germline positions and assigns to each genomic segment (characterised by the same normalised coverage) its respective copy number (cn_i, between 0-5) and the number of copies of the alternate allele (0-cn_i/2). Variant fraction for likely germline variants (defined as positions reported in dbSNP database ^39^ to have an allele frequency >= 30%, 2.5 million positions) was extracted at these positions using ASEQ tool^40.^.

## Data availability

WGS data from FF breast and ovarian cancer samples and associated patient information is publicly restricted access data and can be downloaded from Wellcome Trust Sanger Institute and the International Cancer Genome Consortium ICGC available at the European Genome-phenome Archive EGA (https://www.ebi.ac.uk/ega/studies/EGAS00001001178). The data generated with clinical samples and used in this study are available from source institution, but restrictions apply to the availability of these data, which were used under license for the current study and so are not publicly available. Requests to access this data should be addressed to L.M or C.G.

## Code availability

The deep learning frameworks used here (TensorFlow resp. Keras) are available at https://www.tensorflow.org/ resp. https://keras.io/. The Python libraries used for computation and plotting of the performance metrics (Pandas, Numpy, SciPy, Scikit-Learn, Lifelines, MatPlotLib and Seaborn) are available under https://pandas.pydata.org/, https://numpy.org/, https://www.scipy.org/, https://scikit-learn.org/stable/, https://github.com/CamDavidsonPilon/lifelines/, https://matplotlib.org/ and https://seaborn.pydata.org/ respectively. GII, is a SOPHiA GENETICS proprietary algorithm and is available as part of SOPHiA GENETICS DDM platform.

## Ethical Approval

The study was approved by the Ethics Commission of the Medical University Vienna (EK Nr:2295/2020). All patients provided informed written consent.

## Author contribution

G.A., T.C., C.P. and Z.X. conceived and planned the study. G.G., T.C., J.B. and C.P. developed GII., A.J., X.B and N.A conceived algorithm for annotation of BRCA samples interpretation, J.B. and H.S-S. implemented the algorithm and analyzed variant calls, M.M. and R. K. generated NGS data under A.W. supervision, G.G., T.C., J.B. analyzed NGS data. M.P., L.W-O., C.G. and L.M. conducted the selection, collection, and characterization of ovarian cancer samples as well as DNA extraction. L.M. and C.G. obtained ethical approval for the use of the clinical samples for this study and provided clinical expertise. C.C., C.P. and Z.X. coordinated the study. A.C.M. wrote the first draft of the manuscript. All authors reviewed and approved the final manuscript.

## Acknowledgments

We thank Jean-Baptiste Mignardot for his help with Figure design. We thank Jacqueline Blank and Barbara Neudert for their contribution in performing the DNA extraction of FFPE samples used in this study.

## Supplementary Data

**Supplementary Figure 1**-Heatmap representation of the normalized coverage represented in bins of 100 Kbp lpWGS data for 274 randomly selected data augmented samples. Sex chromosomes are excluded from this analysis.

**Supplementary Figure 2**-**(A)** Heatmap representation of the normalized coverage represented in bins of 100 Kbp lpWGS data for 66 randomly selected data augmented samples. Sex chromosomes are excluded from this analysis. **(B)** Normalized coverage for Clinical-4 sample.

**Supplementary Table S1-** Properties of the clinical samples used in this study.

