## Supplementary figures and images for "A deep learning approach for improved detection of homologous recombination deficiency from shallow genomic profiles"

### Supplementary Figure 1

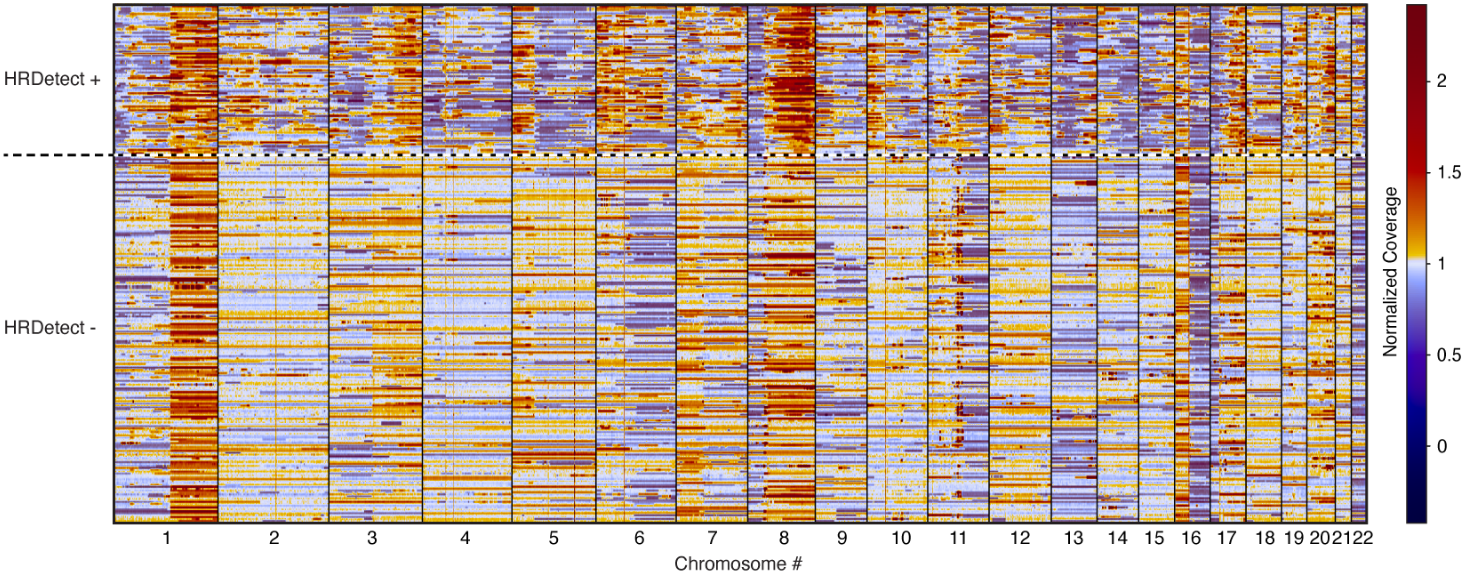

### Supplementary Figure 2

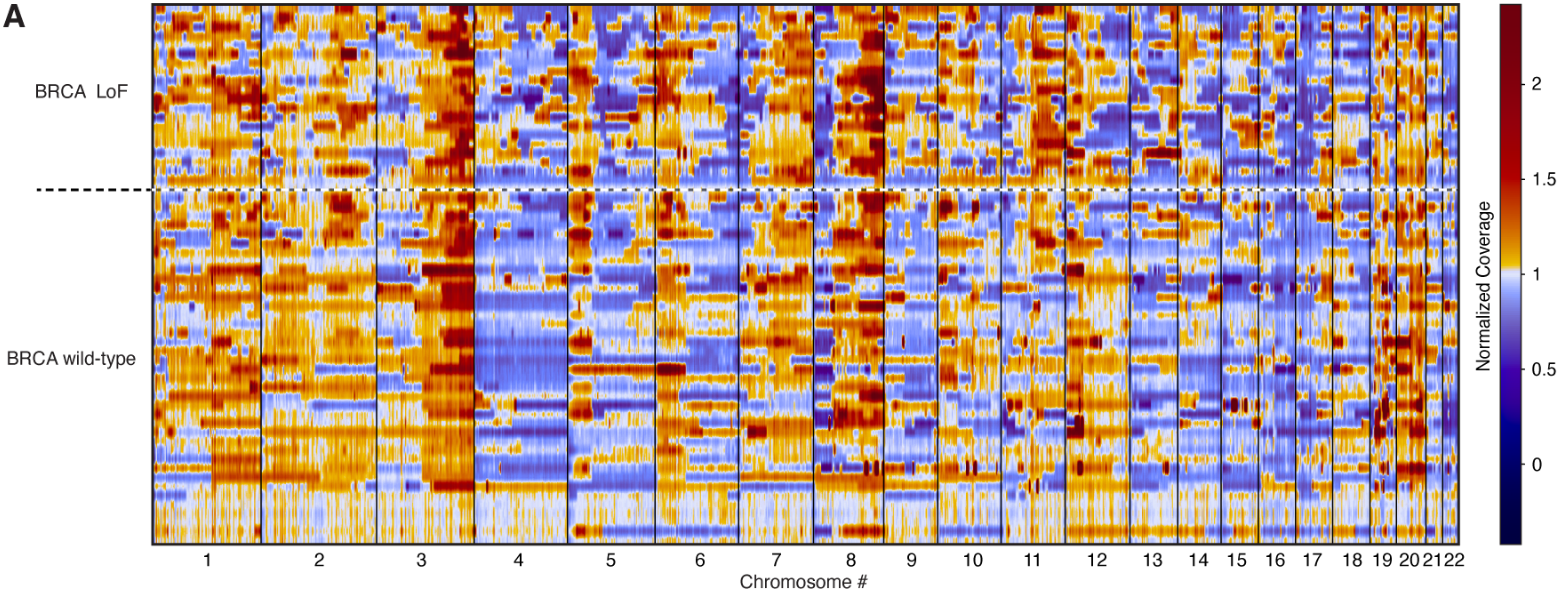

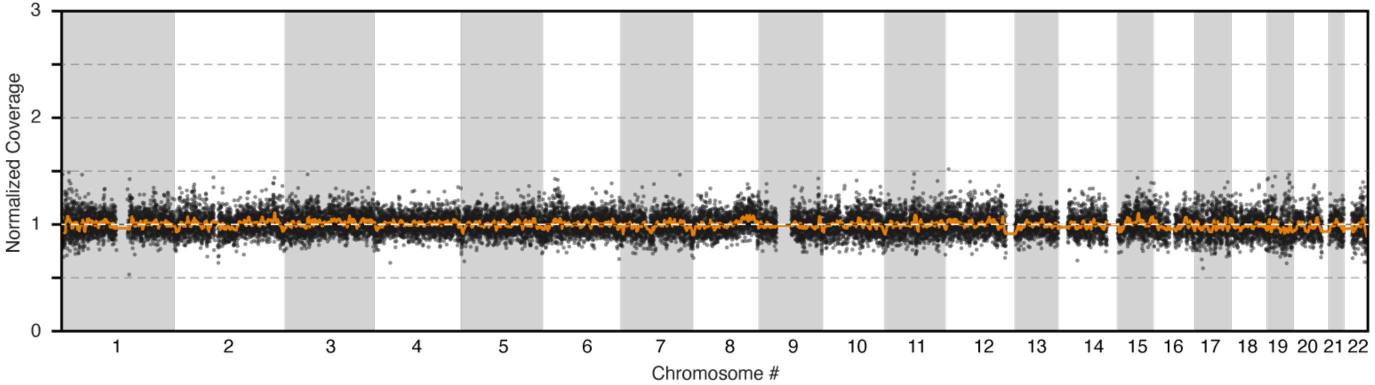
